# Jellyfish blooms - an overlooked hotspot and potential vector for the transmission of antimicrobial resistance in marine environments

**DOI:** 10.1101/2024.07.07.602378

**Authors:** Alan X. Elena, Neža Orel, Peiju Fang, Gerhard J. Herndl, Thomas U. Berendonk, Tinkara Tinta, Uli Klümper

## Abstract

Jellyfish, and gelatinous zooplankton (GZ) in general, represent an important component of marine food webs. Certain GZ species are capable of generating massive blooms of severe environmental impact. These blooms are often followed by a sudden collapse of the entire population, introducing considerable amounts of organic matter (GZ-OM) in the ocean’s interior. GZ-OM represents an abundant substrate to promote bacterial growth and copious colonizable surface for microbial interactions. Hence we hypothesized that this GZ-OM serves as a yet overlooked hotspot for transmitting antimicrobial resistance genes (ARGs) in marine environments. For this we experimentally evolved and analyzed marine microbial communities in microcosms in presence and absence of OM from scyphozoan *Aurelia aurita* s.l. and ctenophore *Mnemiopsis leidyi*. Communities evolved under GZ-OM exposure displayed an up to 4-fold increase in relative ARG and an up to 10-fold increase in abundance of horizontally transferable mobile genetic elements (MGEs) per 16S rRNA gene copy compared to the controls. This trait was consistent across ARG and MGE classes and independent of the GZ species, suggesting that the underlying mechanism is indeed based on the general influx of nutrients and colonizable surfaces. Potential ARG carriers included known key GZ-OM degraders, but also genera containing potential pathogens hinting towards an increased risk of ARG transfer to pathogenic strains. Here, *Vibrio* were pinpointed as potential key species directly associated with several significantly elevated ARGs and MGEs. Subsequent whole-genome sequencing of a *Vibrio* isolate from the microcosm experiment revealed the genetic potential for the mobilization and transfer of ARGs in GZ-OM degrading microbial consortia. With this study, we established the first link between two emerging issues of marine coastal zones, jellyfish blooms and AMR spread, both likely increasing in projected future ocean scenarios.

## 1. Introduction

Gelatinous zooplankton represents an important component of marine food webs inhabiting tropical to polar marine ecosystems. There they represent ∼30% of total biovolume, corresponding to 8-9% of the globally stored carbon in planktonic communities^1^. The most common groups among marine gelatinous zooplankton include ctenophores (comb jellies), medusae (jellyfish), salps, and Chaetognatha. Within the context of climate change, given future ocean projections and considering the adaptability of gelatinous zooplankton to a wide range of environmental conditions these organisms will likely increasingly dominate planktonic marine ecosystems, leading to significant changes in the ocean’s carbon cycle^1–3^. An increase in gelatinous zooplankton abundance, threatening marine ecosystem health and services, has already been recorded worldwide, particularly in anthropogenically impacted coastal areas^2,4^. Due to a combination of life history traits and low metabolic requirements certain gelatinous zooplankton species (e.g., *Mnemiopsis leidyi, Aurelia aurita*)^5,6^ are capable of generating massive blooms, representing an important perturbation to marine ecosystems^7–9^. These blooms are often followed by a sudden collapse of the entire population when a large influx of gelatinous zooplankton detrital organic matter (GZ-OM) distorts the ambient seawater organic matter pool by releasing considerable amounts of bloom-specific particulate, dissolved organic and inorganic matter compounds^10–12^. GZ-OM can then be degraded and consumed at different rates in a cascade by specific microbial assemblages, dominated by copiotrophic bacterial lineages, with consistent metabolic fingerprints^7,9,13^.

In this study, we hypothesize that the GZ-OM introduced into ocean ecosystems upon the decay of bloom events serves as a yet overlooked hotspot for transmitting antimicrobial resistance genes (ARGs) in marine environments.

Antimicrobial resistance (AMR) and the spread of ARGs is one of the major global health challenges^14^ and understanding the biotic and abiotic drivers underlying this spread is crucial to create targeted intervention measures^15,16^. With abundances of ARGs increasing in all ecosystems due to anthropogenic activities^17,18^, marine ecosystems, and their microbiomes are no exception^19,20^. In particular, coastal zones as a likely entry point of ARG-carrying microbes from, for example, wastewater effluents to the marine environments are in the spotlight. Yet, potential links between these two emerging issues of anthropogenically impacted coastal zones, bloom-forming gelatinous zooplankton species, and ARG have not yet been explored. One of the most important ecological mechanisms underlying this spread of ARGs is the conjugative transfer of ARG-encoding mobile genetic elements (MGEs) such as plasmids^21,22^. These conjugative plasmids can not only spread between closely related bacteria but also be transferred to phylogenetically distant bacterial groups^23–25^. Plasmid transfer rates in aquatic environments are particularly elevated when bacterial abundance and activity are high, ensuring high bacterial encounter rates such as in biofilm formed on the surfaces of particles^26–28^. Both these factors are given during the decay of jellyfish bloom events as the released GZ-OM represents an abundant substrate to promote bacterial growth and copious colonizable surfaces for interactions. Furthermore, microbial degraders of gelatinous zooplankton detritus, potentially enriched in AMR, could hitchhike on these organic particle surfaces by drifting with ocean currents over long distances into the oceanic interior^29,30^ and/or connect the microbiomes of upper and bottom ocean layers upon sinking to the ocean floor^31,32^. These bloom events could provide not only a yet overlooked hot spot for the spread of AMR but also a potential vector of AMR/ARG transmission in marine environments.

Consequently, to address the hypothesis that GZ-OM degrading microbial communities provide a hot spot for the enrichment of AMR, we re-analyzed existing metagenomic datasets^7,13^ from microcosm experiments previously conducted and performed and analyzed new, time-resolved, and replicated GZ-OM degradation experiments. This provided insights into how the microbial degradation of biomass from different bloom-forming gelatinous zooplankton species affects the abundance, diversity, and dynamics of AMR and MGEs.

## 2. Material & Methodology

### 2.1. Re-analysed metagenomic datasets

The initial metagenomic datasets used in this study originate from previously conducted experiments on the degradation of gelatinous zooplankton-derived organic matter (hereinafter GZ-OM) described in detail in Tinta *et al.* (2020, 2023) and Fadeev *et al.* (2024)^7,12,13^. Briefly, we conducted short-term microcosm evolution experiments to simulate the scenario potentially experienced by the coastal pelagic microbiome after the decay of GZ blooms. The first set of experiments was conducted using biomass of the cosmopolitan scyphozoan jellyfish, *Aurelia aurita* s.l.^13^. The second set of experiments was conducted using biomass of the lobed ctenophore *Mnemiopsis leidyi*, one of the most notorious marine invasive species^7^. Both species form massive aggregations in several ecosystems around the world, causing a threat to marine ecosystem services. For each evolution experiment we filled six 10 L borosilicate glass microcosms with 0.2 µm filtered aged seawater (ASW), which was inoculated with a 1.2 µm prefiltered coastal microbial community collected in near-surface waters of the northern Adriatic Sea in a ratio of ASW:bacterial inoculum of 9:1. Based on the average abundance of gelatinous zooplankton per m^3^ during typical bloom conditions in our study coastal ecosystem - the northern Adriatic Sea - three experimental microcosms representing the GZ-OM treatment, received 100 mg L^-1^ of GZ-OM (either *A. aurita* or *M. leidyi*). Three microcosms with no GZ-OM amendment served as the control treatment. All microcosms were incubated in the dark at *in situ* temperature (∼ 24 °C) and mixed gently before sampling. Bacterial community dynamics (abundance, production, single-cell metabolic activities) and organic/inorganic nutrients are described in detail in Tinta *et al.* (*A. aurita* dataset)^13^ and Fadeev *et al.* (*M. leidyi* dataset)^7^. At the same time, the microbial inoculum and microbial communities at the peak of bacterial abundance (∼24 hours) were sampled for microbial metagenome and metaproteome (endo- and exo-fraction) analysis from each of the replicate microcosms per treatment. DNA was extracted according to Angel *et al*.^33^ with minor modifications for GZ-OM as described in Tinta *et al.*^12^. Extracted DNA from each replicate was pooled in equimolar amounts before metagenomic sequencing, resulting in one metagenome dataset per treatment. This approach provided early insights into metabolic networks operated by microbial degraders of gelatinous zooplankton organic matter^7,13^ and was here re-analyzed for AMR and MGEs.

### 2.2 Time-resolved and replicated experiment of *M. leidyi* GZ-OM degradation

To complement the already available datasets, we performed a replicated experimental design of the *M. leidyi*-OM microcosms as described in Fadeev *et al.*^7^ and above. To increase the statistical validity of the insights gained, this time we produced metagenomes of the microbial inoculum, as well as each of the triplicate microbial communities in either the presence or absence of *M. leidyi* GZ-OM after 43 and 67 hours.

### 2.3 DNA extraction and Metagenomic sequencing

Metagenomic DNA library preparation, and sequencing were performed as previously described ^12,13^. Raw reads were deposited at NCBI under the accession number PRJNA633735. The obtained raw reads were assessed quality-wise using FastQC^34^. Downstream quality processing was done using BBTools^35^, specifically, quality trimming was done using bbduk, employing a Phred based algorithm and a cut-off of 25 and right and left trimming approach. For the detection of potential phiX phage residual contamination, bbduk was used using a Kmer size of 31 and the integrated phiX fasta sequence.

### 2.4 Metagenomic analysis

The taxonomic composition of each sample was assessed using Kraken2^36^, a k-mer-based approach. For this, we used the Kraken2 PlusPF database and adjusted the confidence to 0.175, a value optimised by using a set of high-quality mock communities (SRR8073716^37^ and SRP436666^38^). To account for the known underestimation of taxonomic assignments, abundance re-classification through a naïve Bayesian approach using Bracken was carried out^39^. Antimicrobial resistance genes and the 16S rRNA gene content in the metagenomes were predicted in each clean metagenomic dataset using the ARGs-OAP V2.0^40^ in which a specific ARG database (Structured Antibiotic Resistance Genes - SARG) constructed from RefSeq ARG hidden Markov Models is used as a reference. Metagenomics reads are aligned to the reference using BLASTx and a 90% identity cutoff. The abundance of reads matching a certain ARG is recalculated taking into consideration the RefSeq sequence length, therefore accounting for uneven coverage and potential over estimation. The 16S rRNA gene content is similarly estimated and then used to normalise the ARG abundance^40^. For mobile genetic element (IS and replicons) relative abundance estimation, a similar approach was used, in these cases, the reference databases were downloaded from ISFinder^41^ and PlasmidFinder^42^ and formatted accordingly using ARGs-OAP make-db command.

### 2.5 *Vibrio splendidus* isolation and whole genome analysis

The *Vibrio splendidus* isolate whose whole genome sequence was analyzed in this study was isolated from an enrichment experiment with *Aurelia aurita* s.l., described in detail in Tinta *et al.*, 2012^13^. For bacterial isolation, 100 µL from GZ-OM microcosms, were spread on modified ZoBell solid agar media and incubated in the dark at 2 °C by gently agitating for 48 h. Single colonies were clean streaked once, inoculated into ZoBell liquid medium, and incubated in the dark at 21°C for 24 h. Bacterial genomic DNA from the resulting colony was extracted immediately with a modified Chelex-based procedure, amplified with universal primers 27F and 1492R, and sent for Sanger sequencing at Macrogen Inc. Furthermore, the bacterial isolate was stored at the culture collection of the Marine Biology Station Piran, Slovenia (in 30% glycerol at − 80°C). After identification as *Vibrio splendidus*, the isolate was regrown from cryo-preserved stock on ZoBell agar plates in the dark at 24°C for 72 h. Then a single colony was inoculated into 6 mL of ZoBell liquid medium, and incubated at room temperature in the dark on a shaker. Four 1 mL replicates of the liquid culture were pelleted by centrifugation at 4000x g for 3 min and bacterial pellets were shipped on dry ice to the sequencing facility (Microsynth AG, Balgach, Switzerland) where high molecular weight DNA was extracted. The DNA was then sequenced using the long-read MinION ONT (Oxford Nanopore Technologies, Oxford, United Kingdom) technique and complemented by short-read paired-end (2 × 75 bp) sequencing on Illumina NextSeq (Illumina, San Diego, CA, USA). Raw long and short reads were quality trimmed using Filtlong V0.2.1 and Trimmomatic V0.39^43^ and used to generate a high-quality hybrid assembly with Unicycler V0.4.8^44^. A total of 3 contigs were assembled and annotated using the RAST server^45^. Antimicrobial resistance genes, replicon-associated structures, and virulence factors were screened using the ResFinder, PlasmidFinder, and VirulenceFactor databases^42,46,47^.

### 2.6. Statistical assessment

Statistical evaluations were carried out in R V4.2.2^48^. Differences in ARG, MGE or bacterial group abundances between sample types were assessed using t-tests with Bonferroni correction for multiple testing. Consistency of effects on ARG abundances across antibiotic classes between samples was assessed through a Wilcoxon signed-rank test with antibiotic classes as the signed variable. Dynamics of ARG and MGE abundances over time were assessed using Pearson correlation tests. For analysis of microbial community and ARG diversity, predicted OTU abundances were transformed using the Hellinger method and Euclidean distances between samples were calculated using the Vegan package in R^49^. Euclidean distances based on relative ARG abundance between each sample were calculated without previous transformation using the Vegan package in R^49^. The correlation between OTU and ARG ordinations was calculated using a symmetric Procrustes approach with ordination plots created using the ggplot2 package^50^. Statistical differences between different groups of samples in the plots were calculated using the analysis of molecular variance (AMOVA) test^51^. Throughout, statistical significance was assigned at p < 0.05. To explore potential ARG-host relationships, pairwise correlations analysis between the ARG and bacterial genera abundances based on Spearman rank correlation were assessed. Only positive correlations showing coefficients ρ > 0.75 were considered significant at p < 0.05 after Benjamini-Hochberg correction for multiple testing. The correlations were displayed through network analysis, performed using the picante package^52^ based on only the significant Spearman rank correlations. The network was visualized using the open-source software Gephi v8.2^53^.

## 3. Results

### 3.1 Antimicrobial resistance dynamics during microbial degradation of gelatinous zooplankton organic matter

To explore whether gelatinous zooplankton organic matter (hereinafter GZ-OM) following the collapse of the gelatinous zooplankton bloom (hereinafter GZ-blooms) could result in the proliferation of antimicrobial resistance (AMR), we first reanalyzed metagenomes from a lab-based microcosm degradation experiments of *Mnemiopsis leidyi*^13^ and *Aurelia aurita* s.l.^7^.

Microcosms were seeded with a coastal marine microbial community in the presence and absence of *M. leidyi* or *A. aurita* s.l. OM and were terminated after the microbial community reached the late exponential growth phase.

In the *M. leidyi* experiment the total relative abundance of antimicrobial resistance genes (ARGs) decreased in the absence of GZ-OM from 1.61 × 10^-2^ ARGs per copy of the 16S rRNA gene in the coastal marine microbiome to 8.18 × 10^-3^ ARGs/16S (Figure 1A). In contrast, the microbiome exposed to GZ-OM was highly enriched in ARGs with a final relative abundance of 5.46 × 10^-2^ ARGs/16S within less than 2 days of exposure (Figure 1A). The main prevalent (>10^-3^ ARGs/16S) antibiotic classes these ARGs conferred resistance to were polymyxins, tetracyclines, quinolones, trimethoprim, fosfomycin, and chloramphenicol (Figure 1B). The observed trend for total ARGs was also consistent at the antibiotic class level: For the majority of antibiotic classes detected ARGs conferring resistance to that class were consistently decreased compared to the original coastal microbiome with the control treatment based on a Wilcoxon signed-rank test with antibiotic classes as the signed variable (p = 0.012, z = -2.49, W = 3, n = 10). Similarly, for the GZ-OM treatment, a significant increase in relative ARG abundance across antibiotic classes was observed when compared to either the original community (p = 0.023, z = -2.23, W = 5, n = 10) or the control treatment (p = 0.037, z = -2.09, W = 7, n = 10).

**Figure 1:**
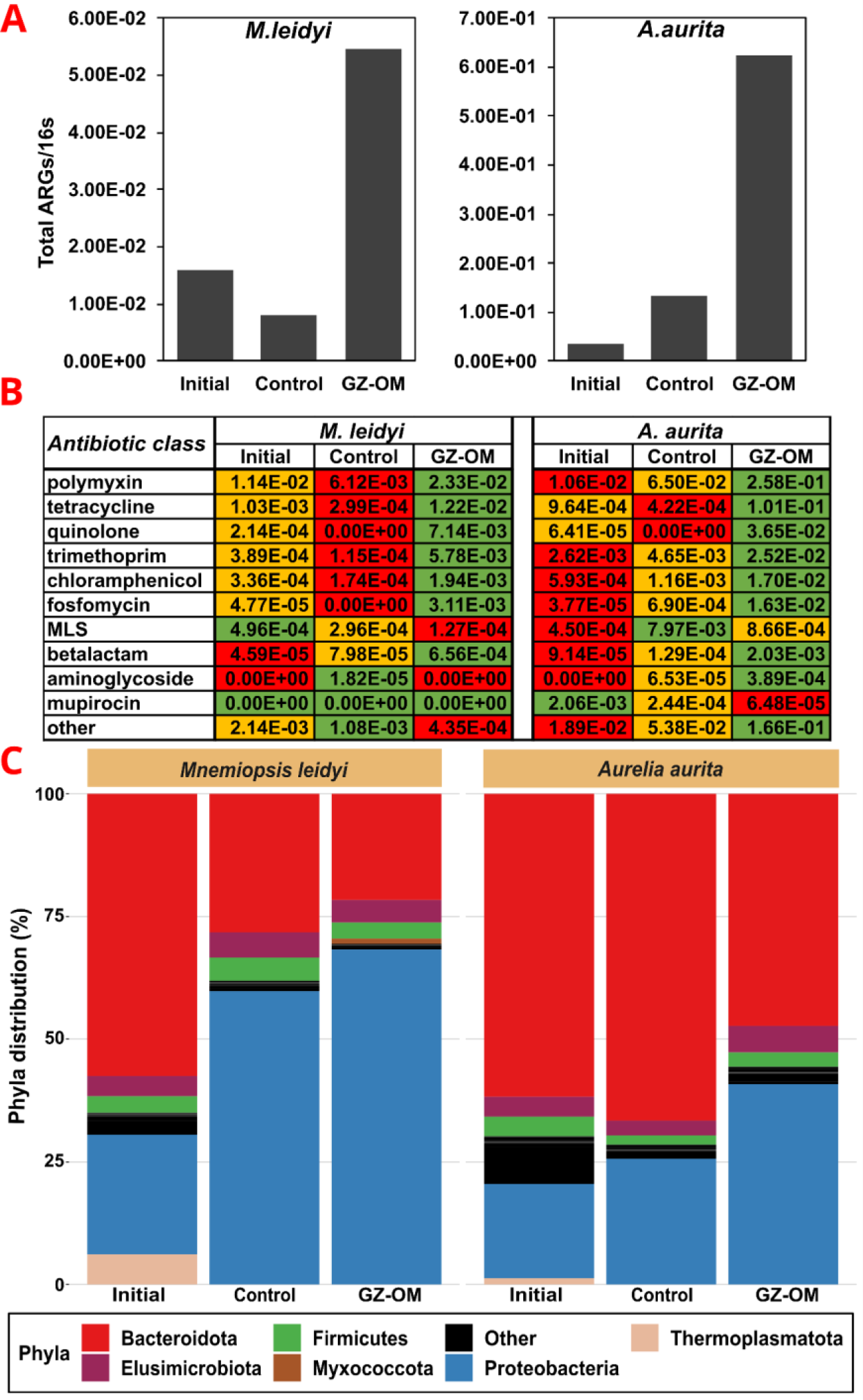
Characterization of the resistome and the microbiome of the initial microbial community as well as those from microcosm experiments in the presence and absence of *Mnemiopsis leidyi or Aurelia aurita GZ- OM*. A: Relative abundance of total ARGs; B: Relative abundance of ARGs based on antibiotic classes they confer resistance to, color-coded for highest (green), intermediate (yellow), and lowest (red) abundance in the initial, control, and jellyfish community. C: Microbial community composition in the microcosm on the phylum level, phyla with abundance below 1% are grouped as others

Similar trends were observed in the *A. aurelia* experiments, where a strong increase in the total relative abundance of ARGs from 3.64 × 10^-2^ in the initial microbiome to 6.23 × 10^-1^ ARGs/16S in the presence of jellyfish biomass was observed. This accounted for a >4-fold increase compared to the no jellyfish control 1.34 × 10^-1^ ARGs/16S (Figure 1A). The observed trend remained again consistent at the antibiotic class level with similar antibiotic classes (polymyxins, tetracyclines, quinolones, trimethoprim, chloramphenicol, and fosfomycin) dominating the resistome: For the majority of ARG classes relative abundances were consistently increased in the jellyfish treatment compared to the initial microbiome (p = 0.0096, z = -2.5887, W = 6, n = 12, Wilcoxon Signed- Rank Test) and the control treatment (p = 0.0232, z = -2.2749, W = 10, n = 12, Figure 1B). In summary, we observed early trends of AMR increasing during GZ-OM degradation independent of the GZ-OM origin species and across antibiotic classes.

### 3.2 Microbial players responsible for GZ-OM degradation and involved in the proliferation of AMR

We next identified the main bacterial groups responsible for GZ-OM degradation and hence likely involved in the observed increase in AMR in the degrading communities. In both experiments (*M. leidyi* & *A. aurita*) the initial microbial communities were mainly dominated by *Bacteroidota* and *Pseudomonadota.* However, a clear shift towards a higher abundance of *Pseudomonadota* at the expense of *Bacteroidota* was observed in both the control as well as the GZ-OM treatments (Figure 1C). This shift was far more pronounced in those microcosms exposed to either of the two types of GZ-OM (Figure 1C). We identified candidate genera that might have contributed to the observed increase in AMR as they displayed a high abundance (>0.1% rel. abundance) and were found at >5-fold increased abundance in the GZ-OM compared to the control treatment. For both types of GZ-OM, these included well-known GZ-OM degraders (*Pseudoalteromonas, Vibrio, Alteromonas*)^7,13^, but also genera that contain human or animal pathogens *(Enterobacter, Escherichia-Shigella, Vibrio, Pajaroellobacter, Francisella*)^54^. In addition, *Oleiphilus, Anaerosinus*, *Acinetobacter,* and *Arsenophonus* increased in abundance in the *M. leidyi* experiment. In summary, similar microbial genera dominated both *M. leidyi* and *A. aurita* GZ-OM degradation, indicating that despite the limited number of replicates, the observed trends are generalizable for biomass degradation of diverse GZ species. Still, as in these original studies we only operate with a single metagenome dataset per treatment and at a single time point, insights gained remained limited. Thus, an additional GZ-OM degradation experiment with replicates and samples taken over different time points was performed and analyzed.

### 3.3 Antimicrobial resistance genes are consistently enriched in microbial communities degrading *Mnemiopsis leidyi* GZ-OM

To gain statistically valid insights whether degradation of GZ-OM indeed increases AMR proliferation in the marine microbiome, we analyzed metagenomes from the newly performed *M. leidyi*-OM degradation experiment with the appropriate replication, sampling, and sequencing at three time points representing the microbial inoculum (T0h), the microbial community at the late exponential phase of their growth (T43h) and the microbial community after reaching stationary growth phase (T67h). Again, total ARG relative abundance increased in the GZ-OM treatment from 0.91 × 10^-2^ (T0h) to 3.85 ± 1.85 × 10^-2^ (T67h) ARGs per 16S rRNA gene with a significant increase rate of 4.9 × 10^-4^ ARGs/16S per hour (R = 0.68, p = 0.048, Pearson correlation, Figure 2A).

**Figure 2:**
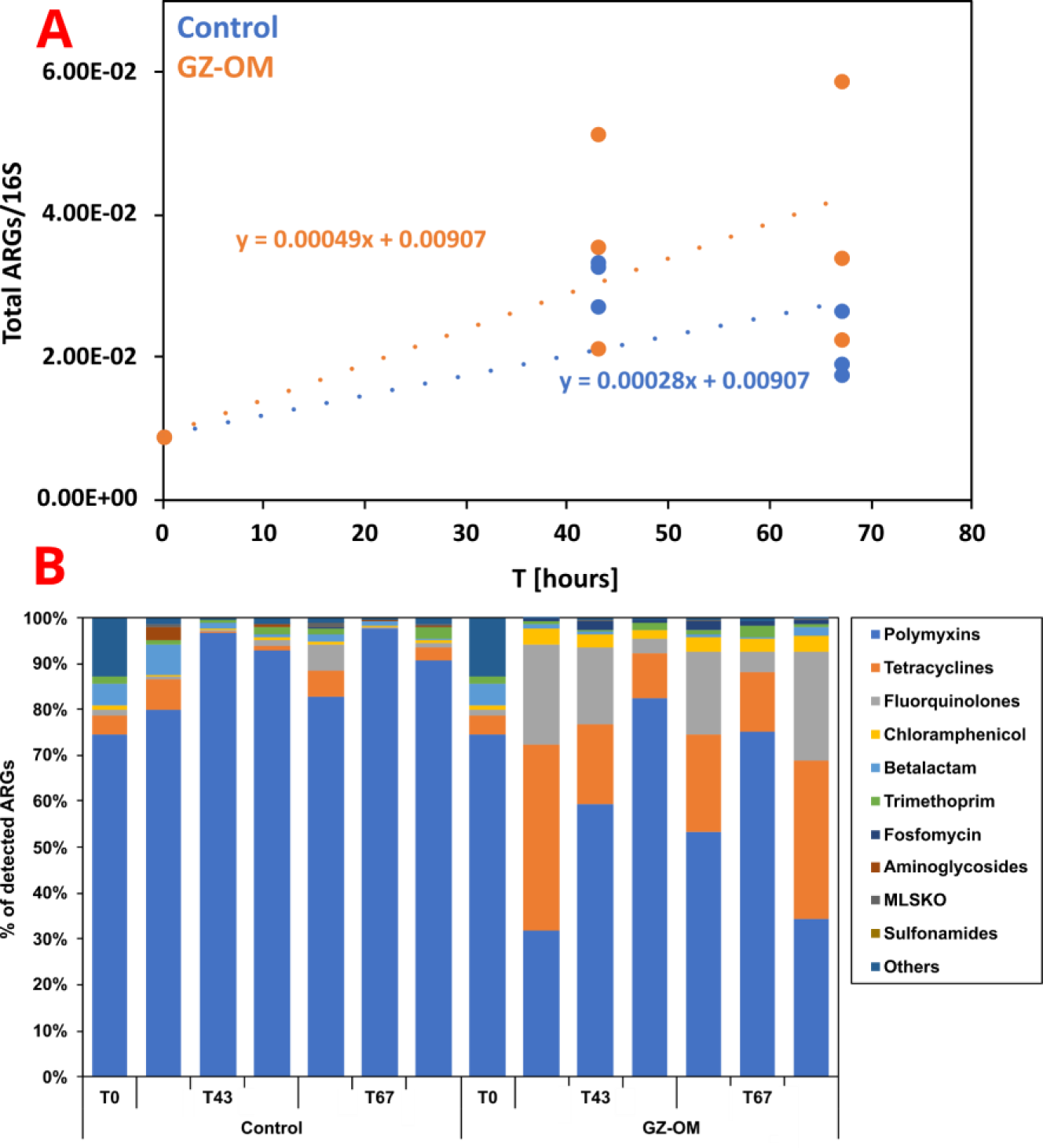
Resistome dynamics in the *M. leidyi* GZ-OM and control microcosms over time. A: Pearson correlation of total ARGs per 16S rRNA gene in the metagenomes over time. B: Proportion of antibiotic resistance gene classes in the total ARGs of the metagenomes (MLSKO = macrolide, lincosamide, streptogramin, ketolide and oxazolidinone)

However, no significant difference in ARG levels over time (m = 2.8 × 10^-4^ ARGs/16S per hour, R = 0.65, p = 0.121, Figure 2A) was detected in the control treatment without GZ-OM addition. This increase at the ARG class level displayed a clear effect of GZ-OM especially for tetracycline and fluoroquinolone ARGs. Tetracycline ARGs significantly increased in proportion from initially 4.1% of the total ARGs to 20.0 ± 13.2% in GZ-OM treatments (p < 0.01, t-test). In contrast, the proportion of tetracycline ARGs remained rather stable in the control treatment (2.8 ± 2.8%, p > 0.05, t-test) (Figure 2B). Similarly, fluoroquinolone resistance genes increased from 1.1% to 12.7 ± 9.5% in GZ-OM treatments (p<0.01, t-test) while remaining stable in the control at 1.5 ± 2.1% (p > 0.05, t-test) (Figure 2B). This increase in proportion was mirrored in the relative abundances increasing by more than one order of magnitude from initially 3.79 × 10^-4^ to 9.87 ± 8.45 × 10^-3^ tetracycline ARGs/16S (p < 0.01, t-test) and from 9.98 × 10^-5^ to 6.48 ± 5.29 × 10^-3^ fluoroquinolone ARGs/16S (p < 0.01, t-test). For the remaining antibiotic classes, the shifts were smaller and remained below one order of magnitude in relative abundance.

### 3.4 Mobile genetic elements are consistently enriched in GZ-OM degrading microbial communities

As increases in AMR are of particular risk if associated with a rise in MGEs that can facilitate their transfer to pathogenic strains, we similarly analyzed the MGE content of the metagenomic samples with a particular focus on insertion sequences (IS). While the number of detected IS families remained consistent between 18 and 23 independent of any treatment, the relative IS abundance per 16S rRNA gene mirrored that of ARGs. In both initial datasets IS relative abundance was increased in the GZ-OM treatment (*M. leidyi*: 3.41 × 10^-1^ IS/16S; *A. aurita*: 2.83 × 10^-1^ IS/16S; n=1) compared to their respective control (*M. leidyi*: 1.97 × 10^-2^ IS/16S; *A. aurita*: 1.98 × 10^-2^ IS/16S; n=1) (Figure 3A). Similar in the time-resolved experiment a statistically significant increase in IS relative abundance was observed for the GZ-OM treatment at both timepoints (43h: 1.21 ± 0.96 × 10^-1^ vs. 3.67 ± 2.89 × 10^-2^ IS/16S; 67h: 1.72 ± 1.40 × 10^-1^ vs. 2.82 ± 1.91 × 10^-2^ IS/16S; both p <0.05, n=3, t-test; Figure 3A). The observed effect was largely consistent across the different individual IS families, however, a particular significant increase in the proportion of the IS*110* family in the *M. leidyi*-OM (28.6 ± 6.7%) compared to the control treatment (8.1 ± 2.4%; p < 0.001, t-test) stood out (Figure 3b). Within this IS*110* family, we detected 36 individual IS out of which IS*Visp2*, IS*Visp6*, IS*Spi5*, IS*Ptu2*, IS*Spi2* and IS*Cps8* were significantly enriched in the M. leidyi-OM treatment. Similarly in the *A. aurita*-OM treatment, we observed 27 individual IS belonging to the IS*110* family. Again, IS*Visp6*, IS*Visp2*, IS*Ptu2*, IS*Spi5* and IS*Cps8* were almost exclusively associated with GZ-OM. Members of the IS*110* family have been reported to form ARG carrying transposons^55^, highlighting the relevance of the increased presence of these IS associated with GZ-OM degrading communities. In general, IS*Visp2* and IS*Visp6*, native to *Vibrio splendidus*, seemed to be the most abundant of the IS*110* members detected both in *M. leiydi*- and *A. aurita*-OM treatments. This places *Vibrio* not only as a relevant GZ-OM degrader but also as a potential genus involved in horizontal ARG dissemination.

**Figure 3:**
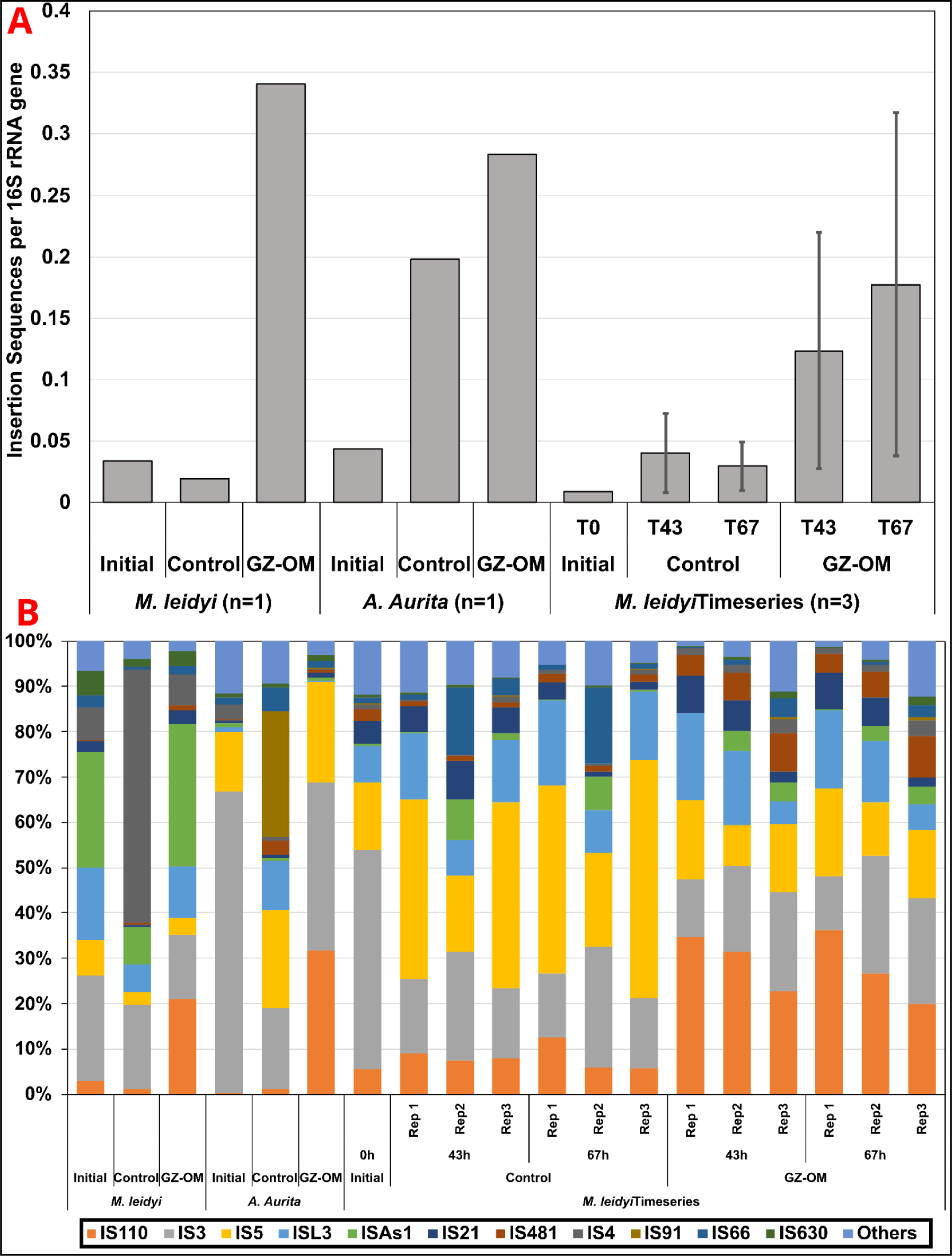
Mobile genetic element dynamics in the GZ-OM and control microcosm metagenomes. A: Relative abundance of insertion sequences per 16S rRNA gene B: Proportion of IS families among the total IS content in the metagenomes

### 3.5 Increase in antimicrobial resistance during *Mnemiopsis leidyi* biomass degradation are related to the microbial community composition

As GZ biomass is unlikely to contain large amounts of selective agents that are responsible for the observed increase in ARGs and MGEs, a more likely explanation is a shift in microbial community composition during the degradation process. Indeed, for both the GZ-OM as well as the control treatment, a clear shift in community composition at the genus level was observed when comparing the microcosm communities to the initial community composition (p < 0.05, AMOVA, Figure 3A). However, GZ-OM treatment communities consistently grouped further apart from the initial community (0.627 ± 0.029 average Hellinger transformed taxonomic distance) than the control treatment (0.534 ± 0.069, p=0.011, ANOVA, Figure 3A). Moreover, the control and GZ treatments were grouped apart from each other (p<0.05, AMOVA). Still, no clear effect of the sampling time between T43h and T67h could be observed for either the GZ-OM or control samples (all p>0.05), indicating that the shifts in community composition appeared early during GZ-OM degradation. The individual ARG diversity between GZ-OM-, control treatment, and initial community was again significantly different (all p < 0.05, AMOVA, Figure 3B). Phylogenetic diversity explained a significant proportion of the observed ARG diversity using Procrustes analysis (Correlation index = 0.8019; sum of squares = 0.366, p = 0.0001). Consequently, we aimed to identify those bacteria responsible for this shift in the resistome.

**Figure 3:**
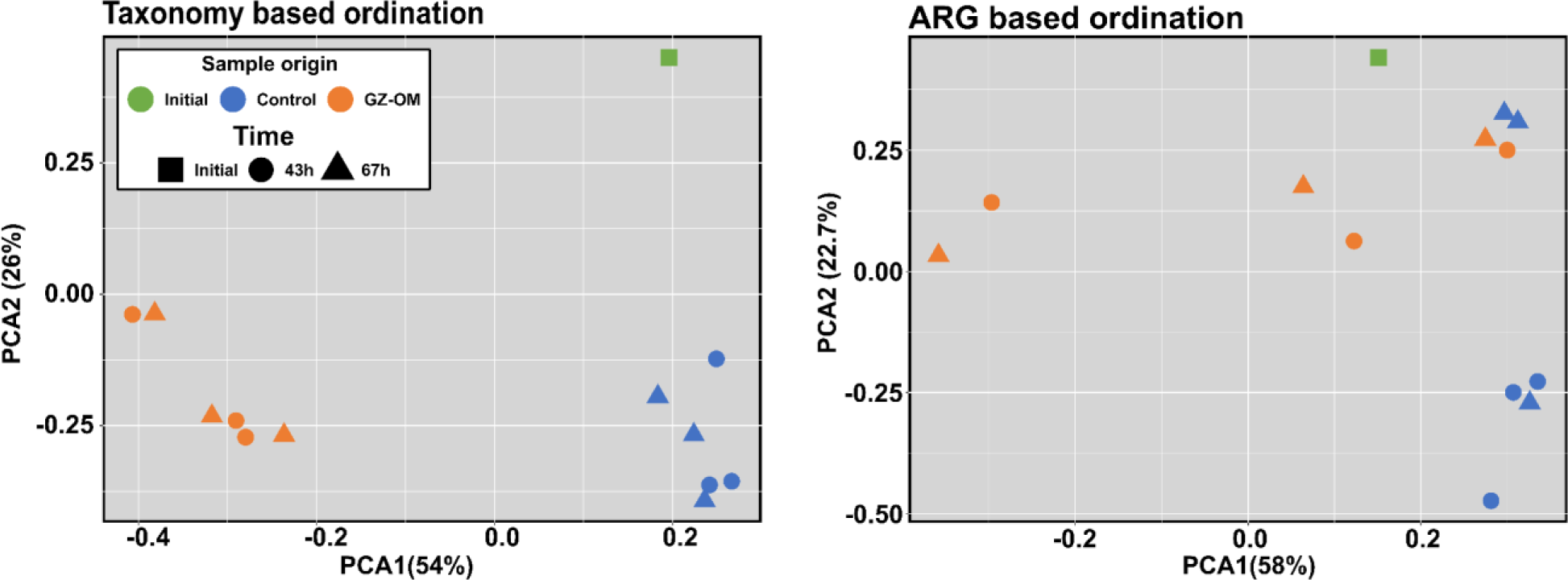
Microbial community and ARG dynamics in the microcosm experiments: A) PCA plot of genus-level microbial community composition based on Euclidean distances after Hellinger transformation of the data; B) PCA plot of individual ARG-level resistome diversity of the microbiomes based on Euclidean distances.

### 3.6 Bacterial genera associated with GZ-OM degradation including human and animal pathogens are potential ARG hosts

To identify which bacteria are responsible for the observed increase in ARGs in the GZ-OM treatment we analyzed the microbial community compositions. On the phylum level, most notably, in both treatments, the abundance of the most prevalent *Pseudomonadota* significantly increased compared to the initial community from 69.38% to 79.92 ± 3.97% (control) and 81.67 ± 2.99% (jellyfish) (both p < 0.05, t-test, Figure 4A), but no significant differences between the GZ-OM and the control treatment were observed. For other phyla, no clear significant trends were detected. However, since genus level diversity analysis suggested that differences between the control and the GZ-OM treatment exist (Figure 3A), we identified those genera that were highly abundant in general (>0.5% rel. abundance), significantly elevated compared to the initial community (all p < 0.05, t-test) and found at >5-fold significantly increased abundance in the GZ-OM compared to the control treatment (all p < 0.05, ANOVA). These probably contributed to the observed increase in ARG levels. The majority of the identified GZ-OM treatment-associated genera belonged to the phylum *Pseudomonadota*. Among these, the most prevalent were *Vibrio* (4.07 ± 3.09%), *Pseudoalteromonas* (3.13 ± 0.60%), and *Thalassotalea* (2.11 ± 0.09%), all previously identified as capable GZ-OM degraders^7,13^. Additionally, two typical marine genera were identified: *Colwellia* (0.69 ± 0.08%), associated with a psychrophilic lifestyle^56^, and *Algicola* (0.53 ± 0.25%), a regular colonizer of algal surfaces^57^. Still, similar to the preliminary dataset analyzed, genera regularly hosting human pathogens^54^ such as *Enterobacter* (1.37 ± 0.35%) or the aforementioned *Vibrio* and animal pathogens^58,59^ such as *Arsenophonus* (1.13 ± 0.29) and *Pajaroellobacter* (0.63 ± 0.26%) were identified as GZ-OM associated.

**Figure 4:**
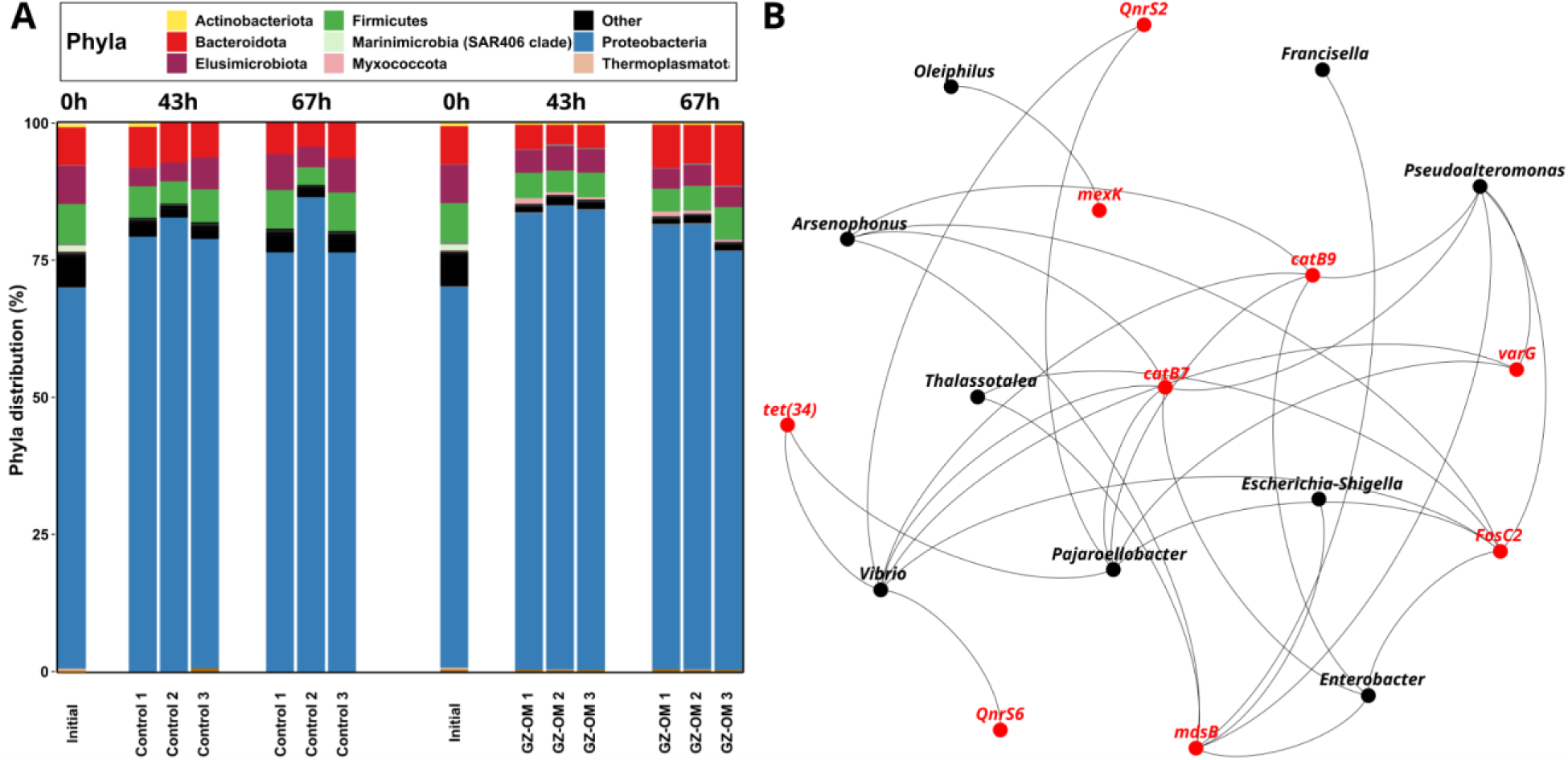
Microbial community composition and potential ARG hosts in the GZ-OM degradation microcosm experiments A) Community composition at the phylum level over time. Phyla with abundance below 1% are grouped as others B) Network analysis displaying the correlations in relative abundance between bacterial genera and identified ARGs based on significant (p < 0.05) Spearman correlations with positive correlation coefficients of ρ > 0.75.

To identify whether those GZ-OM degradation-associated genera are indeed linked with ARGs as their potential hosts we performed a network analysis between ARG and genera abundance in the entire dataset revealing a high number of positive genera-genera correlations. Still, when extracting exclusively those 31 connections of ARGs with genera (Figure 4B) it became apparent, that exclusively the 9 previously identified GZ-OM degradation-associated genera could be identified as ARG hosts based on network analysis. Each of these genera had at least one significant connection to an ARG and displayed on average 3.44 ± 2.30 connections to ARGs. Out of the 9 ARGs for which host connections could be inferred, 7 displayed connections to multiple hosts, indicating their potential mobility, which is of concern considering that both marine and human/animal pathogen-associated genera were identified to be GZ-OM-associated. Among the identified host genera, *Vibrio* displayed the highest number of connections with 7 individual ARGs. Moreover, *Vibrio* was the lone potential host of quinolone ARG *qnr*S6, while sharing host association of quinolone ARG *qnr*S2 and tetracycline ARG *tet*(34) with *Pajaroellobacter*. That is particularly relevant, as we demonstrated that tetracycline and quinolone ARGs displayed the highest increase in GZ-OM treatments.

### 3.7 Analysis of *Vibrio* isolates from the GZ-OM degradation experiment

As previously stated, *Vibrio* spp. has a highly relevant role not only as a degrader, but also as a potential source of ARGs and their horizontal dissemination. We consequently used a *Vibrio* isolate (A06) from the GZ-OM degradation microcosm experiment to analyze whether this highly enriched GZ-OM degradation-associated genus contains ARGs and MGEs that would display the potential of increased horizontal gene transfer of ARGs during the degradation of GZ-OM. Hence, the isolated *Vibrio* strain A06 was sequenced and annotated. For taxonomic characterization, reference quality genomic sequences of 69 *Vibrio* isolates were retrieved from Enterobase^60^, and the 16S rRNA gene sequences extracted and aligned to those of the isolate in a maximum likelihood tree (Figure 5). Eleven distinct clusters, mostly associated with individual species, were observed with isolate A06 identified as belonging to the *V. splendidus* group.

**Figure 5:**
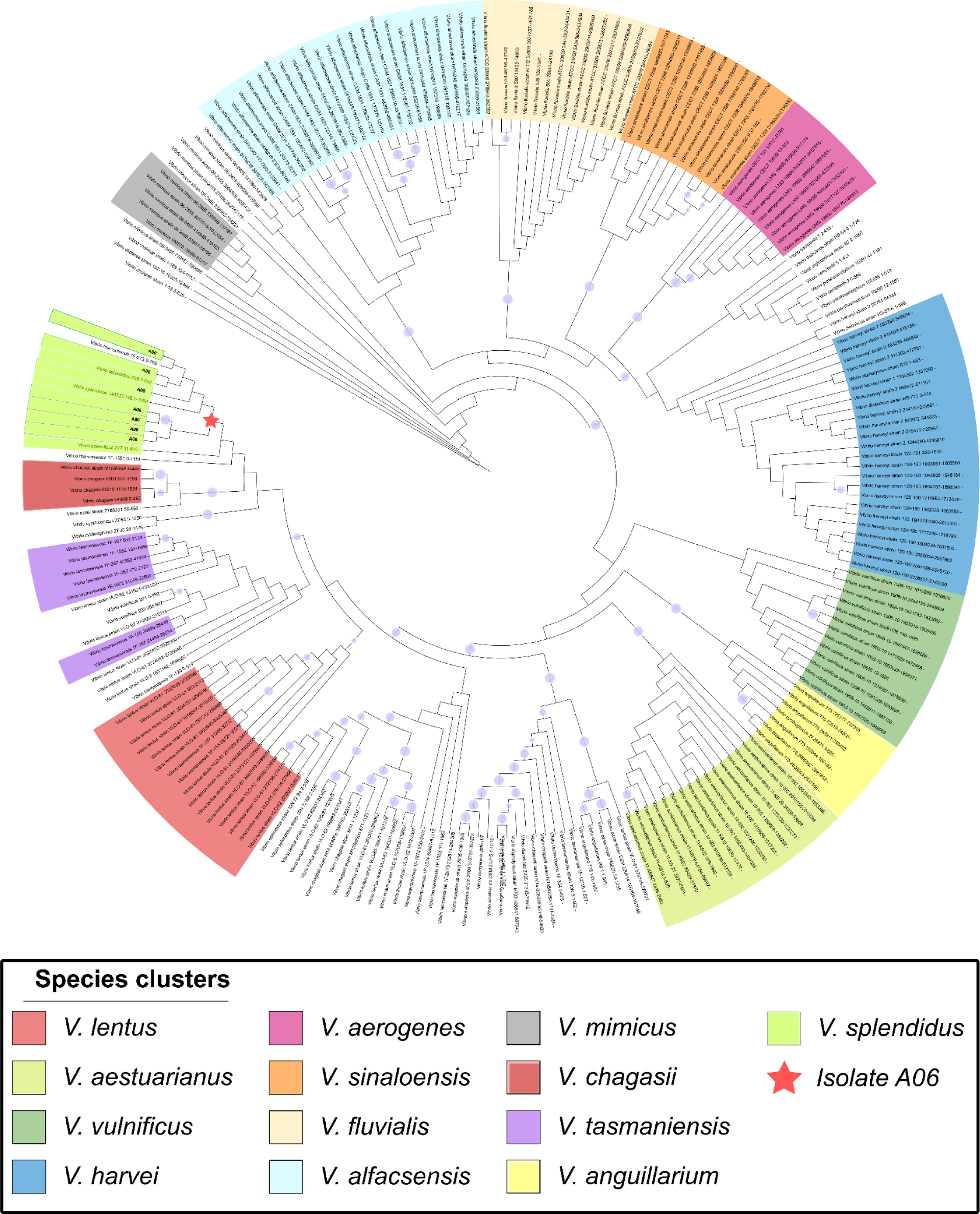
Maximum likelihood tree showing different species within the *Vibrio* genus. Each branch represents an individual copy of the 16S rRNA gene. Colors mark species clusters. Blue circles show bootstrap support values between 0.9 and 1 (related to size).

The assembled *V. splendidus* genome consisted of three circular structures of 3.7, 2.1, and 0.17 Mbps with normalized depths/copy numbers of 1x, 0.98x, and 1.58x. While the first two structures matched several Vibrio species’ first and second chromosomes, the third contained plasmid- related structures. A total of 31 AMR-related genes were distributed across the first two genetic structures (16 in Chromosome 1 and 15 in Chromosome 2), most of which (29/31) coded for multidrug efflux pumps including *mdtL/A/K/C/H*, *mepA*, *emrD/E*, *cat1*, *bmrA,* and *norM*. The two remaining predicted ARGs were the tetracycline resistance gene *tet34* (Chromosome 1) and a putative *qnr* gene (Chromosome 2), typically associated with diminished susceptibility towards fluoroquinolones. The latter was closely related to *qnr*S sequences from RefSeq (Bootstrap > 0.99). Moreover, 38 transposases belonging mostly to IS families *3*, *4*, and *110* were found in chromosome 1. Chromosome 2 harbored 23 additional transposases, the majority of which belonged to IS*4* and IS*110* families. Insertion sequences from these two families have been observed in more than 200 different bacterial and archaeal species^61^ and have been associated with ARG mobility in environmental bacterial communities^62^ and human clinical isolates^63,64^. In the case of A06, no evidence of ARGs or virulence factors within a range of ISs belonging to these families was found. However, several composite transposons harbored hypothetical proteins, suggesting a potentially active role in the incorporation of exogenous genetic material.

The third assembled circular genetic structure was compatible with a plasmid: *repA*, a *tra* region (incomplete), a partition system (*parAB*), and a toxin/antitoxin system (*phd*/*doc*) were identified. A 662 bp origin of replication (*oriC*) was annotated immediately downstream from the predicted *repA* gene. This replicase gene could not be assigned to any incompatibility group present in the PlasmidFinder database, instead, it showed 100% coverage and 98.23% identity with the *repA* of another non-typed plasmid recovered from *Vibrio crassostreae* (AN: AP025478.1). The structure of the plasmid harbored by A06 showed little similarity with the sequences present in the NCBI database with two *Vibrio* genetic structures (AP025478.1 and KP795560.1) being the closest relatives (Figure 6).

**Figure 6:**
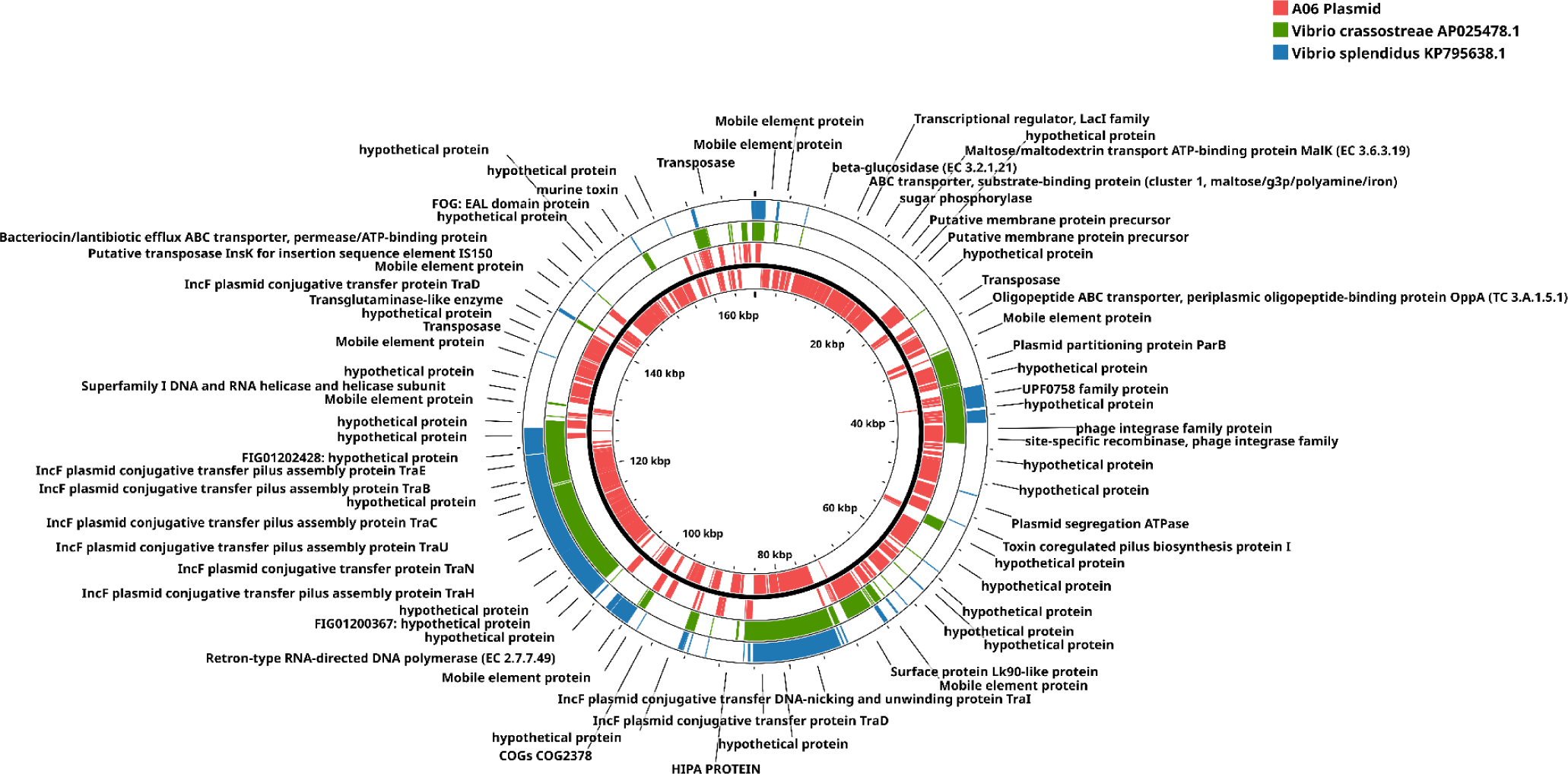
Comparison between the plasmid sequence of A06 (red) and the two closest genetic records in the NCBI database, AP025478.1 (green) and KP795560.1 (blue). The red blocks indicate ORFs in the A06 plasmid while the blue and green blocks represent identical regions in the corresponding genetic structures.

Several ABC-type efflux pumps were detected on the plasmid, most of these related to inorganic toxin efflux. Moreover, the plasmid carried an ABC pump involved in bacteriocin elimination, flanked by two insertion sequences (IS*150* and IS*Sod8*). While not directly related to AMR, the gene *hipA* was detected on the plasmid. The product of this gene (HipA) has been reported to cause reversible “dormancy” of *E. coli*, slowing down cell growth and favoring a persister phenotype that is notoriously tolerant to drugs like β-lactams^65,66^. Similar to the chromosomes, a considerable amount of 22 insertion sequences was detected, the most common families being IS*6* and IS*5*. The former is a well-studied family that has major implications in the clinic and is generally related to the incorporation of promoter sequences that could upregulate flanking genes^67^.

In summary, based on an individual *Vibrio* isolate, we identified the genetic potential for the mobilization and transfer of ARGs through plasmid and IS structures in GZ-OM degrading microbial consortia.

## 4. Discussion

With this study, we established the first link between two emerging issues of marine coastal zones, jellyfish blooms and AMR spread, both likely increasing in projected future ocean scenarios^5,68^. Metagenomic analysis of marine microbial communities exposed to GZ-OM confirmed our hypothesis that decaying jellyfish blooms represent a yet overlooked hot spot of AMR proliferation in marine environments. Already after 2-4 days of exposure to GZ-OM, we recorded an up to 4- fold increase in relative ARG abundance per 16S rRNA gene copy in the degrader communities compared to ambient marine microbiomes. This increase becomes particularly relevant when considering that bacterial production rates due to the nutrient influx through degradable GZ-OM in the otherwise nutrient-poor marine environment can be up to one order of magnitude elevated^5^ with absolute bacterial biomass increasing 10 to 100-fold in the microcosms^7,12^, with equal numbers being reported for natural ecosystems exposed to GZ-OM after bloom events^69^. Combining this increase in absolute bacterial and relative ARG abundances, jellyfish blooms are predicted to result in an absolute increase of ARG abundance by several orders of magnitude compared to the surrounding marine microbiomes.

The observed trait was consistent, independent of the gelatinous zooplankton species and the year of the experiment, suggesting that the underlying mechanism of this increase in AMR is based on the general influx of nutrients and colonizable surfaces through GZ-OM. Still future work should aim at disentangling the individual contributions of these two general mechanisms, which are further supported by the phylogenetic diversity of the colonizing bacterial communities being highly similar across GZ-OM from different species both in our analysis as well as in the literature^70–73^. It furthermore proved to be a significant explanatory variable for the observed ARG diversity and increased ARG abundance. Potential carriers of these increasing ARGs were consistent in all our datasets and included *Pseudoalteromonas*, *Vibrio*, and *Alteromonas*, known key GZ-OM degraders^7,8,13,72,73^, as well as *Thalassotalea*, *Colwellia* associated with psychrophilic lifestyle^56^, *Algicola,* regular colonizers of algal surfaces^57^ and other potential degraders of hydrocarbons in marine environment (*Oleiphilus*, *Anaerosinus*). More importantly, when considering the risks associated with the observed enrichment of ARGs, several genera that contain known potential human or animal pathogens (*Enterobacter*, *Escherichia-Shigella*, *Acinetobacter*, *Vibrio*, *Pajaroellobacter*, *Francisella*, *Arsenophonus*)^54^ were identified to not only be significantly increased in relative abundance in the degrading communities but also correlated with specific ARGs as their potential carriers. This is consistent with previous reports that jellyfish- colonizing microbiomes regularly include elevated proportions of potential human pathogenic strains^72,73^.

The simultaneous increased abundances of potential pathogens and ARGs in the degrading communities do on their own not immediately translate into an elevated risk if ARGs are not transferred. Our data provide strong indication that such horizontal acquisition of ARGs by these potential pathogenic strains is indeed taking place. First, similar to ARGs, the relative abundance of MGEs in the GZ-OM degrading communities was significantly elevated. These ARG-encoding MGEs have the potential to be transferred even to phylogenetically distant bacterial groups^23–25^ and horizontal gene transfer rates are particularly elevated when bacterial abundances and activity are high and bacteria have high encounter rates^26–28^ such as in biofilms formed on GZ- OM particles. The observed high connectivity between marine environmental and potentially pathogenic species in the ARG-genera co-occurrence network as co-hosts of specific identical ARGs suggests that this scenario indeed occurs. Second, genomic analysis of the *A. aurita*-associated *V. splendidus* strain revealed a high number of insertion sequences, and numerous multidrug-efflux pumps but also the ARGs *tet*(34) and *qnrS* carried in the two identified chromosomes. The quinolone resistance encoding *qnr* genes are generally plasmid-associated^74^, while *tet*(34) has been observed in a broad range of environmental hosts^75^, suggesting its general mobility. In addition, a yet unknown plasmidic structure was identified that hosts several insertion sequences, a copy of the *hipA* gene able to induce a dormant state that favors the persistence of ARGs^65,66^, and a bacteriocin efflux pump as part of a compound transposon. Together, this provides strong indication that the GZ-OM degrading communities have members that indeed possess the necessary genomic plasticity that provides a high potential of acting as donors and recipients of horizontally transferable ARGs. This could constitute a significant risk, as these GZ- OM colonizing communities enriched in AMR and potential pathogens that could acquire novel ARGs can hitchhike on these particle surfaces by drifting with ocean currents over long distances in the ocean interior and coastal environments where exposure to higher marine organisms (e.g., also commercially important species) and/or humans is likely^29,30^.

When considering gelatinous zooplankton detritus as a hotspot for the marine spread of AMR, it is also likely that living gelatinous zooplankton colonized by bacteria could equally play a role. Here it is additionally relevant to consider their life stage-specific features. During the polyp stage, meroplanktonic gelatinous zooplankton species are mostly found in coastal areas that frequently are highly anthropogenically impacted (e.g., pillars of industrial ports), where they can accumulate different types of pollutants^76–78^. These could provide a (co-)selective potential for ARGs of their microbial colonizers^79,80^, while also being in more direct contact with potential colonizers enriched in ARGs (e.g., from wastewater discharged into the ocean^81^). After strobillation, the ephyrae develop to the adult medusa stage, which can drift and/or swim with ocean currents over long distances, and in this way represent an overlooked route of AMR (and ARG) to otherwise not impacted environments especially since some GZ, like *Mnemiopsis leidyi*, are invasive species. During this stage, as they are efficient grazers of a significant part of the ocean’s planktonic production^2,82^ they can further accumulate nanoparticles, microplastic debris^83–86^, heavy metals and pollutants^87^ which can again (co-)select for ARGs and increase horizontal gene transfer rates of the colonizing microbes^24,26,79,88^. These unexplored aspects regarding the spread of AMR in connection with GZ need to be studied and taken into account, especially when considering harvesting GZ for food, fertilizers, medicine, and cosmetics, or considering their use in wastewater treatment applications^89,90^.

## 5. Conclusions

In conclusion, we here provide evidence that jellyfish blooms are a quintessential “One Health” issue where decreasing environmental health is immediately connected to benign effects on human health by amplifying the spread of antimicrobial resistance genes and their potential transfer to human pathogens. This is of particular relevance as both issues are likely to increase in importance with current climate change projections. While detailed research into the underlying mechanisms remains necessary, our initial study already clearly demonstrates that improving ocean health remains imperative not only to control the occurrence of future jellyfish blooms but also to mitigate the spread of antimicrobial resistance. Potential mitigation measures include the control of overfishing to maintain natural jellyfish predators, management of climate change impacts, and most importantly reducing nutrient pollution through the discharge of waste waters into the oceans. This practice not only promotes jellyfish blooms but also leads to the introduction of pathogenic bacteria able to engage in horizontal ARG uptake as a consequence of jellyfish blooms.

## 6. Data Availability

The datasets supporting the conclusions of this article are included within the article and its additional files or available through the corresponding authors upon reasonable request. Original sequencing data is available in the NCBI sequencing read archive under project accession numbers PRJEB63998 (*M. leidyi* experiments), PRJNA633735 (*A. aurita* experiments).

## Acknowledgments/Funding

We acknowledge Eduard Fadeev for conducting preliminary analysis on some of the metagenomic datasets used in the study. UK & TUB were supported by the Explore-AMR and the JPIAMR SEARCHER project funded by the Bundesministerium für Bildung und Forschung under grant numbers 01DO2200 & 01KI2404A. AXE, UK & TUB were supported by the ACRAS-R project funded by Bundesministerium für Bildung und Forschung under grant number 16GW0355. PF was supported through the China Scholarship Council (CSC) under grant number 202004910327. This project received funding from the European Union’s Horizon 2020 Research and Innovation Program under the Marie Skłodowska-Curie Grant Agreement No. 793778. GJH was funded by the Austrian Science Fund (FWF) project I04978. TT was supported by the Slovenian Research Agency under grant number ARRS J7-2599 and by the Slovenian Research Agency (Research Core Funding No. P1-0237). Responsibility for the information and views expressed in the manuscript lies entirely with the authors.

## 7. Competing Interests

The authors declare no competing interests.

## 8. CRediT Author Contributions

**Alan X. Elena:** Conceptualization, Methodology, Validation, Formal analysis, Investigation, Data Curation, Writing - Original Draft, Writing - Review & Editing, Visualization

**Neža Orel:** Validation, Formal analysis, Data Curation, Writing - Review & Editing

**Peiju Fang:** Methodology, Formal analysis, Investigation, Writing - Review & Editing, Visualization, Funding acquisition

**Gerhard J. Herndl:** Resources, Writing - Review & Editing, Supervision, Project administration, Funding acquisition

**Thomas U. Berendonk:** Resources, Writing - Review & Editing, Supervision, Project administration, Funding acquisition

**Tinkara Tinta:** Conceptualization, Methodology, Validation, Formal analysis, Investigation, Resources, Data Curation, Writing - Original Draft, Writing - Review & Editing, Visualization, Supervision, Project administration, Funding acquisition

**Uli Klümper:** Conceptualization, Methodology, Validation, Formal analysis, Resources, Data Curation, Writing - Original Draft, Writing - Review & Editing, Visualization, Supervision, Project administration, Funding acquisition

## Notes

### Competing Interest Statement

The authors have declared no competing interest.

